# Chitinase 3-like-1 is a Therapeutic Target That Mediates the Effects of Aging in COVID-19

**DOI:** 10.1101/2021.01.05.425478

**Authors:** Suchitra Kamle, Bing Ma, Chuan Hua He, Bedia Akosman, Yang Zhou, Chang Min Lee, Wafik S. El-Deiry, Kelsey Huntington, Olin Liang, Jason T. Machan, Min-Jong Kang, Hyeon Jun Shin, Emiko Mizoguchi, Chun Geun Lee, Jack A. Elias

## Abstract

COVID-19 is caused by the SARS-CoV-2 (SC2) virus and is more prevalent and severe in the elderly and patients with comorbid diseases (CM). Because chitinase 3-like-1 (CHI3L1) is induced during aging and CM, the relationships between CHI3L1 and SC2 were investigated. Here we demonstrate that CHI3L1 is a potent stimulator of the SC2 receptor ACE2 and viral spike protein priming proteases (SPP), that ACE2 and SPP are induced during aging and that anti-CHI3L1, kasugamycin and inhibitors of phosphorylation, abrogate these ACE2- and SPP-inductive events. Human studies also demonstrated that the levels of circulating CHI3L1 are increased in the elderly and patients with CM where they correlate with COVID-19 severity. These studies demonstrate that CHI3L1 is a potent stimulator of ACE2 and SPP; that this induction is a major mechanism contributing to the effects of aging during SC2 infection and that CHI3L1 coopts the CHI3L1 axis to augment SC2 infection. CHI3L1 plays a critical role in the pathogenesis of and is an attractive therapeutic target in COVID-19.

## Introduction

SARS-CoV-2 (SC2) is a novel coronavirus that was initially appreciated in man in 2019. It is highly transmissible and has caused a global pandemic killing >1.5 million people and infecting > 80 million people worldwide as of December, 2020 (1-7). The disease caused by SC2, COVID-19, was initially noted to manifest as a pneumonia with fever, cough, fatigue and dyspnea as major manifestations (8). However, it is now known to be a systemic disease with manifestations in many organs (9, 10). In addition, it is now known that there is a spectrum of disease severity with patients that are asymptomatic, 10-20% of patients that require hospitalization (4, 5, 11). and patient with respiratory failure that require intensive care (12). Unique clinical features have also been described including pulmonary fibrosis, an increased frequency of vascular thrombotic events, a coagulopathy and pulmonary angiitis (13, 14). However, the cellular and molecular events that account for this impressive clinical and pathologic heterogeneity are poorly understood.

One of the most unique features of COVID-19 is its impressive relationship to aging and comorbid disorders. The elderly is at increased risk of contracting COVID-19 and experience higher complication and case fatality rates (15, 16). COVID-19 has also been devastating effects on and elderly and residents of congregate care facilities (17, 18). Similarly, approximately 50% of hospitalized COVID-19 patients have preexisting medical conditions including diabetes, hypertension, obesity and metabolic syndrome, cardiovascular disease and chronic lung diseases like COPD and asthma. These co-morbidities associate with enhanced susceptibility to SC2 infection, more severe disease and a higher mortality (1-3, 5, 15, 19, 20). Surprisingly, the cellular and molecular events that underlie the effects of aging and comorbid diseases in COVID-19 have not been defined.

SC2 interacts with cells via its spike (S) protein which binds to its host cell receptor angiotensin converting enzyme 2 (ACE2) which mediates viral entry (21-24). After binding, the S protein is processed by the S priming proteases (SPP) TMPRSS2, Cathepsin L (CTSL) and in some cases FURIN into S1 and S2 subunits. The latter mediates the fusion of the viral envelope and cell plasma membrane which is required for virus-cell entry (15, 19, 23). In the human disease, SC2 passes through the mucus membranes of the upper and lower respiratory tracts and infects epithelial cells via this ACE2-S protein-protease mechanism (15, 19, 25). The virus can then enter the bloodstream and, via this viremia, infect other cells and organs that express ACE2 and SPP (26). It has been proposed that the levels of expression of ACE2 and the SPP play key roles in determining the magnitude and organ location of the infection and the severity of the disease (15, 19, 27). However, the mechanisms that SC2 uses to generate its effects in different tissues have not been defined. Furthermore, although ACE2 is expressed widely in human tissues (15), the critical processes that regulate ACE2 expression and activation have only recently begun to be investigated.

Virus infection of mammalian cells leads to innate and adaptive immune responses that restrict viral replication, augment viral clearance and limit tissue damage and disease severity (28). To allow infection to occur many viruses have developed strategies to evade and or suppress these immune responses (28-30). They can also co-opt host responses to augment viral replication (28-30). These events can drastically influence disease pathogenesis, the course of the infection, disease severity and viral persistence in the host (28-30). Compared to other positive stranded (+) RNA viruses, coronaviruses have exceptionally large genomes and employ complex genome expression strategies (28). Many of the coronavirus genes participate in virus-host interplay to create an optimal environment for viral replication (28). However, the host pathways that SC2 co-opts to foster infection and replication have not been adequately defined and the possibility that SC2 utilization of host responses contributes to the mechanisms by which aging and comorbid diseases augment the prevalence and severity of COVID-19 has not been addressed.

Chitinase 3-like-1 (CHI3L1; also called YKL-40 in human and Chil1 or BRP-39 in mouse) is an evolutionarily conserved member of the 18 glycosyl hydrolase (GH 18) gene family that is produced by a spectrum of cells in response to a variety of injury and cytokine stimuli (31-35). It is the cornerstone of a critical pathway that is activated during injury and inflammation, regulates innate and adaptive immunity and heals and protects (32, 34, 36-38). The latter is mediated by its ability to inhibit apoptosis and other forms of cell death while driving fibroproliferative repair (36, 38). CHI3L1 is readily detected in the circulation of normals and the levels of circulating and tissue CHI3L1 are increased in diseases characterized by inflammation, injury, remodeling and repair (31-33, 37, 39-46). Interestingly CHI3L1 is also expressed in an exaggerated manner in aging and in the same comorbid diseases that are risk factors for COVID-19 (47-54). CHI3L1 and ACE2 are also both mediators of pulmonary protective responses (36, 55-58). In light of these impressive similarities we hypothesized that the induction of ACE2 and SPP is part of the CHI3L1-induced healing and repair response. We also hypothesized that SC2 co-opts the host CHI3L1 axis to augment its ability to infect, spread and generate disease. To address these hypotheses, we used genetically modified mice, *in vitro* approaches and human cohorts. These studies demonstrated that (a) CHI3L1 is a potent stimulator of ACE2 and SPP in pulmonary epithelial cells and vascular cells; (b) that the expression of ACE2 and SPP are increased during murine aging and that these aging-induced inductive events are mediated by CHI3L1; and (c) that interventions that alter CHI3L1 effector responses are also potent inhibitors of ACE2 and SPP and viral infection. Lastly, human studies demonstrated that the levels of circulating CHI3L1 are increased in COVID-19 (+) patients that are elderly, have comorbid diseases and manifest severe COVID-19 disease. These findings support the concept that CHI3L1 plays a critical role in the pathogenesis of and is an attractive therapeutic target in COVID-19. They also provide a mechanistic explanation for how aging and comorbid diseases contribute to the pathogenesis of COVID-19.

## Results

### CHI3L1 regulation of pulmonary ACE2 and SPP *in vivo*

To begin to address the relationship(s) between CHI3L1 and SC2, we characterized the effects of CHI3L1 on the expression and accumulation of ACE2 and SPP *in vivo*. In these studies, we compared the expression of Ace2 and SPP in lungs from inducible CC10 promoter-driven CHI3L1 overexpressing Tg mice (CHI3L1 Tg) and wild type (WT) controls. These studies revealed the induction of Ace2, Tmprss2 and Ctsl in lungs from CHI3L1 Tg mice (Fig. 1, A and B). Ace2 protein was most prominent in airway epithelium where it colocalized with CC10 (Fig. 1C and Supplemental Fig. S1). Ace2 was also seen in alveoli where it co-localized with pro-SP-C (Fig. 1C and Supplemental Fig. S1). Tmprss2 and Ctsl were also prominently expressed in airway epithelial cells (Fig. 1, C and D and Fig. S1) and TMPRSS2 was also seen in alveolar macrophages (Fig. 1D).

**Fig 1.**
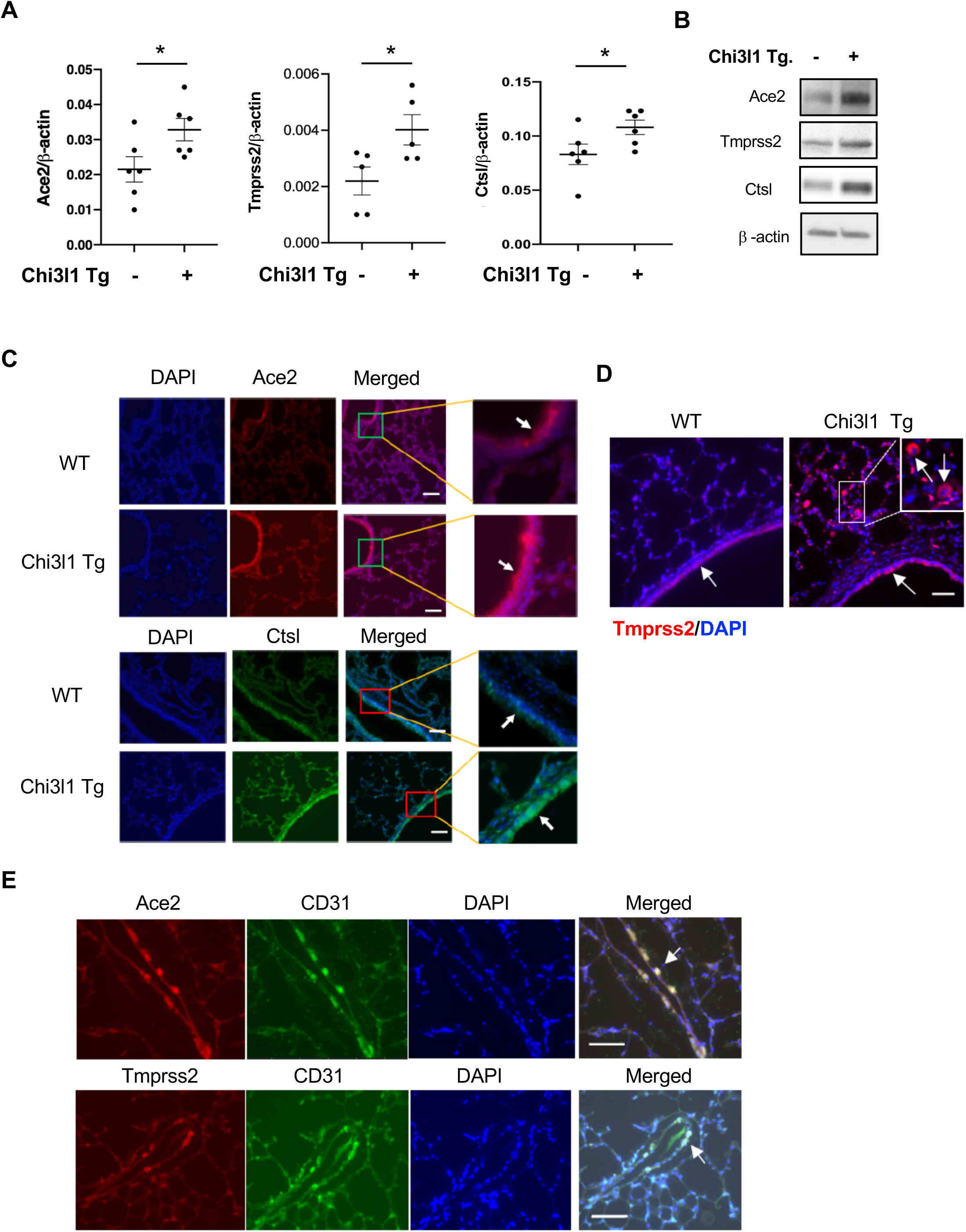
CHI3L1 stimulates pulmonary ACE2 and SPP. 8-week-old WT (-) and Chi3l1 Tg (+) mice were sacrificed after 2 weeks of transgene induction with Doxycycline. The levels of Ace2 and SPP mRNA and protein in the lungs from WT and Chi3l1 Tg mice were evaluated using lung lysates and paraffin tissue blocks from these animals. (A) Comparisons of levels of expression of Ace2 and SPP in lungs from WT (Tg (-)) and Chi3l1 Tg (+) mice using semi-quantitative real-time RT-PCR analysis indexed to β-actin controls. (B) Western immunoblot comparisons of levels of Ace2 and SPP in lungs from WT (Tg-) and Chi3l1 Tg (+) mice. (C-D) Immunohistochemical evaluations of Ace2, Ctsl and Tmprss2 in lungs from WT and Chi3l1 Tg mice. (E) Double label IHC comparing localization of Ace2, Tmprss2 and CD31 in the lungs from Chi3l1 Tg mice. Arrows on panels C-E indicate the stain (+) cells. Each value in panel A is from a different animal and the mean±SEM are illustrated. Panels B-E are representative of at least 3 separate evaluations. Ace2, murine angiotensin converting enzyme 2; Tmprss2, transmembrane serine protease 2; Ctsl, Cathepsin L. Scale bars=100μm. \**P*<0.05 (Student *t*-test).

Tg CHI3L1 also augmented the expression and accumulation of Ace2 and SPP in pulmonary blood vessels. Immunohistochemistry demonstrated that Ace2 and Tmprss2 co-localized with CD-31 on endothelial cells and transgelin (+) vascular smooth muscle cells (Fig. 1E and Supplemental Fig. S1). When viewed in combination these studies demonstrate that CHI3L1 is a potent stimulator of epithelial and vascular cell ACE2 and SPP *in vivo*.

### CHI3L1 regulation of pulmonary ACE2 and SPP *in vitro*

I*n vitro* experiments were next undertaken to determine if rCHI3L1 regulated the expression and or accumulation of ACE2 and SPP. These studies demonstrated that rCHI3L1 stimulated ACE2 and SPP (TMPRSS2, Cathepsin L) gene expression and protein accumulation in human Calu-3 epithelial cells in a time-and dose-dependent manner (Fig. 2, A and B). These effects were not Calu-3 cell specific because similar results were obtained with A549 epithelial cells, primary human small airway epithelial cells (HSAEC cells) and lung fibroblasts (Supplemental Figs. S2 and S3). When viewed in combination, these studies demonstrate that CHI3L1 stimulates ACE2 and SPP in a variety of cells *in vitro*.

**Fig 2.**
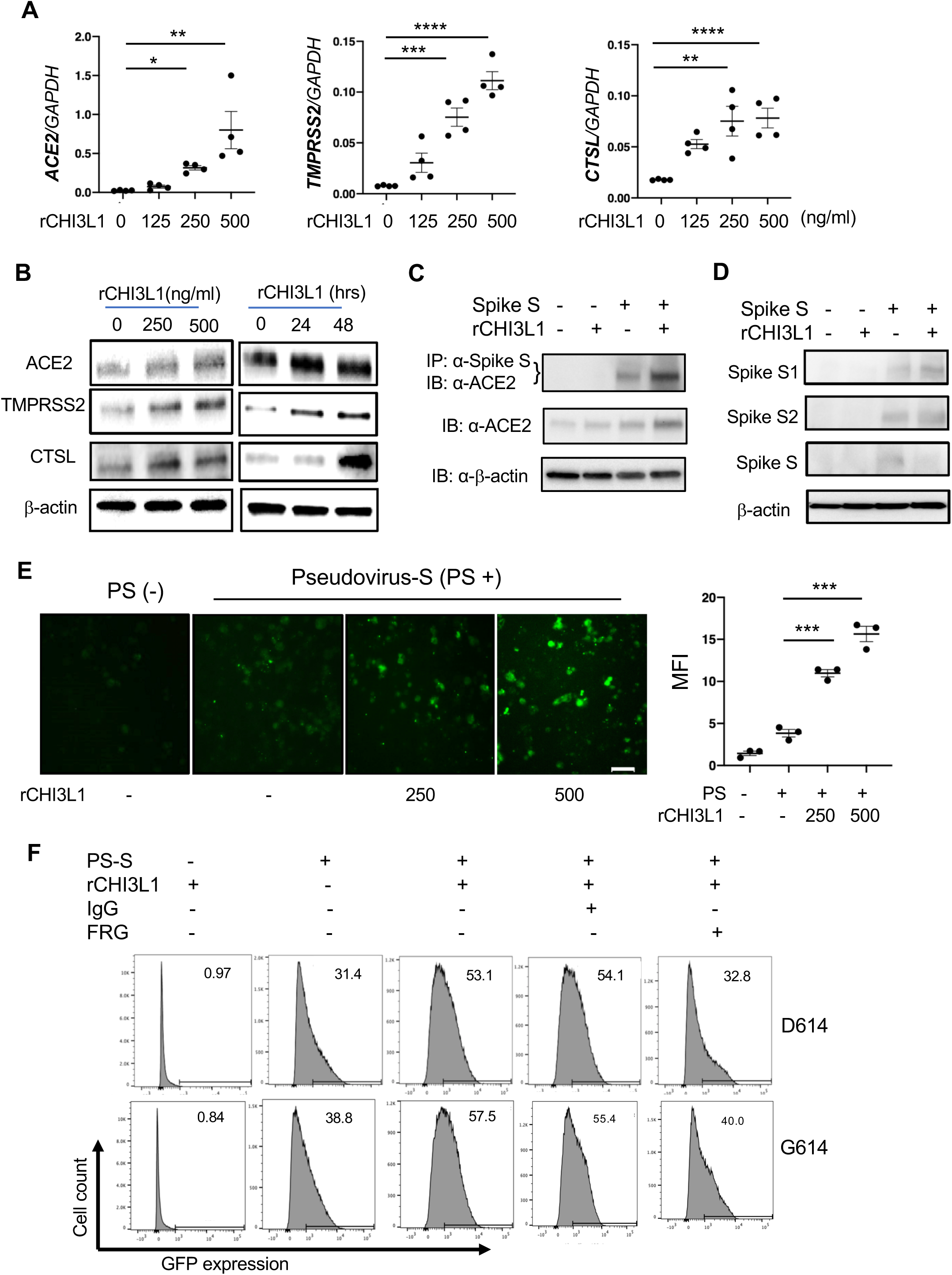
CHI3L1 stimulates ACE2 and SPP *in vitro* and enhances S protein processing and cellular integration. (A) Calu-3 lung epithelial cells were incubated with the noted concentrations of recombinant (r) CHI3L1 (ng/ml) or vehicle control (rCHI3L1=0) for 24 hrs then subjected to RT-PCR to quantitate the levels of mRNA encoding ACE2 and the SPP. (B) Western blot evaluations of the dose response and kinetics of CHI3L1 stimulation of ACE2 and SPP protein accumulation in Calu-3 cells. (C) Calu-3 cells were incubated with vehicle (PBS; CHI3L1 (-)) or rCHI3L1 (500ng/ml) for 24 hrs, recombinant S protein of SARS-COV2 (SC2) was added and the incubation continued for an additional 2hrs. The cells were then harvested, lysates were prepared and co-immunoprecipitation (Co-IP) and immunoblot assays were undertaken. (D) Calu-3 cells were incubated with vehicle (PBS; CHI3L1 (-)) or rCHI3L1 (500ng/ml) for 24 hrs, the cells were harvested, and the recombinant S protein of SARS-COV2 (SC2) was added to the lysates and the incubation continued for an additional 2hrs. The un-cleaved S and cleaved S1 and S2 proteins were evaluated using Western immunoblotting. (E) Calu-3 cells were incubated with vehicle (rCHI3L1(-)) or the noted concentrations of rCHI3L1 for 24 hrs. They were then transfected with a pseudovirus containing the S protein (PS; D614 variant) from SC2 and a GFP expression construct and incubated for additional 24hr then evaluated using fluorescent microscopy. Quantification of mean fluorescent intensity (MFI) can be seen in the dot plot on the right. (F) Calu-3 cells were incubated with rCHI3L1 or vehicle control for 48 hrs in the presence or absence of antibody against CHI3L1 (FRG) or control antibody (IgG). Pseudovirus S (PS; D614 and G614 variants) that expressed GFP was added as noted and GFP expression was evaluated by flow cytometry. Values in panel A and E are mean±SEM. Panels B-F are the representative gel, fluorescent images and FACS data obtained from at least 3 independent experiments. MFI, mean fluorescent intensity. The numbers in the panel F represent % of GFP positive cells. \**P*<0.05, \*\**P*<0.01, \*\*\**P*<0.001, \*\*\*\**P*<0.0001 (One-Way ANOVA with post hoc Dunnett’s multiple comparison test). Scale bar=50μm.

### Consequences of ACE2 and SPP induction

To understand the consequences of CHI3L1 induction of ACE2 and SPP, studies were undertaken to determine if this induction augmented the ACE2 receptor binding of Spike (S) protein and or the metabolism of S into its S1 and S2 subunits. In these experiments lysates were prepared from Calu-3 cells incubated with rCHI3L1 or vehicle control and recombinant (r) spike protein was added. To assess S-receptor binding, immunoprecipitation and immunoblots were serially undertaken with anti-ACE2 and anti-S respectively. To assess S metabolism, the cell lysates were incubated for 24 hours with rS and Western blots were undertaken with S1 and S2 specific antibodies. As can be seen in Figs. 2C and 2D, CHI3L1 stimulation was associated with enhanced ACE2-S binding and enhanced metabolism of S into its respective subunits.

To further address the consequences of these inductive events, Pseudovirus moieties were generated by incorporating the SC2 “S” protein into lentivirus moieties which contained a GFP marker. The infectious capacity of these pseudovirus S moieties was then evaluated using cells incubated with rCHI3L1 or control vehicle. In keeping with the findings noted above, rCHI3L1 induction of ACE2 and SPP augmented pseudovirus S incorporation into lung epithelial cells. (Fig. 2E). FACS evaluations further confirmed these observations by demonstrating that rCHI3L1 enhanced the cellular integration of S-protein containing pseudovirus and that treatment with anti-CHI3L1 monoclonal antibody (FRG) ameliorated rCHI3L1-stimulated pseudovirus cell integration (Fig. 2F). CHI3L1 similarly enhanced the cellular integration of pseudovirus containing S sequence with D416 or G416 and FRG treatment inhibited pseudoviral infection regardless of S sequence (Fig. 2F). When viewed in combination, these studies demonstrate that CHI3L1 stimulation of ACE2 and SPP increases S-ACE2 binding, S metabolism and pseudovirus S infection.

### Monoclonal and small molecule CHI3L1-based therapeutics

The finding that CHI3L1 is a potent stimulator of ACE2 and SPP raises the interesting possibility that CHI3L1 may be a useful therapeutic target in COVID-19. To begin to address this possibility, we characterized the effects of a humanized monoclonal antibody against CHI3L1 developed in our laboratory (called FRG), and a small molecule Chitinase 1 and CHI3L1 inhibitor called kasugamycin *in vitro* and *in vivo*. The monoclonal inhibitor of CHI3L1 (FRG) was a potent inhibitor of CHI3L1 stimulation of epithelial cell ACE2 and SPP *in vitro* (Fig. 3, A and B). FRG was also a powerful inhibitor of Tg CHI3L1 induction of ACE2 and SPP *in vivo* (Fig. 3C). Interestingly, kasugamycin was a similarly powerful inhibitor of CHI3L1 induction of ACE2 and SPP *in vitro* and *in vivo* (Fig. 3, D&E). These studies highlight antibody-based and small molecule CHI3L1 inhibitors that control ACE2 and SPP and have promise as therapeutics in COVID-19.

**Fig 3.**
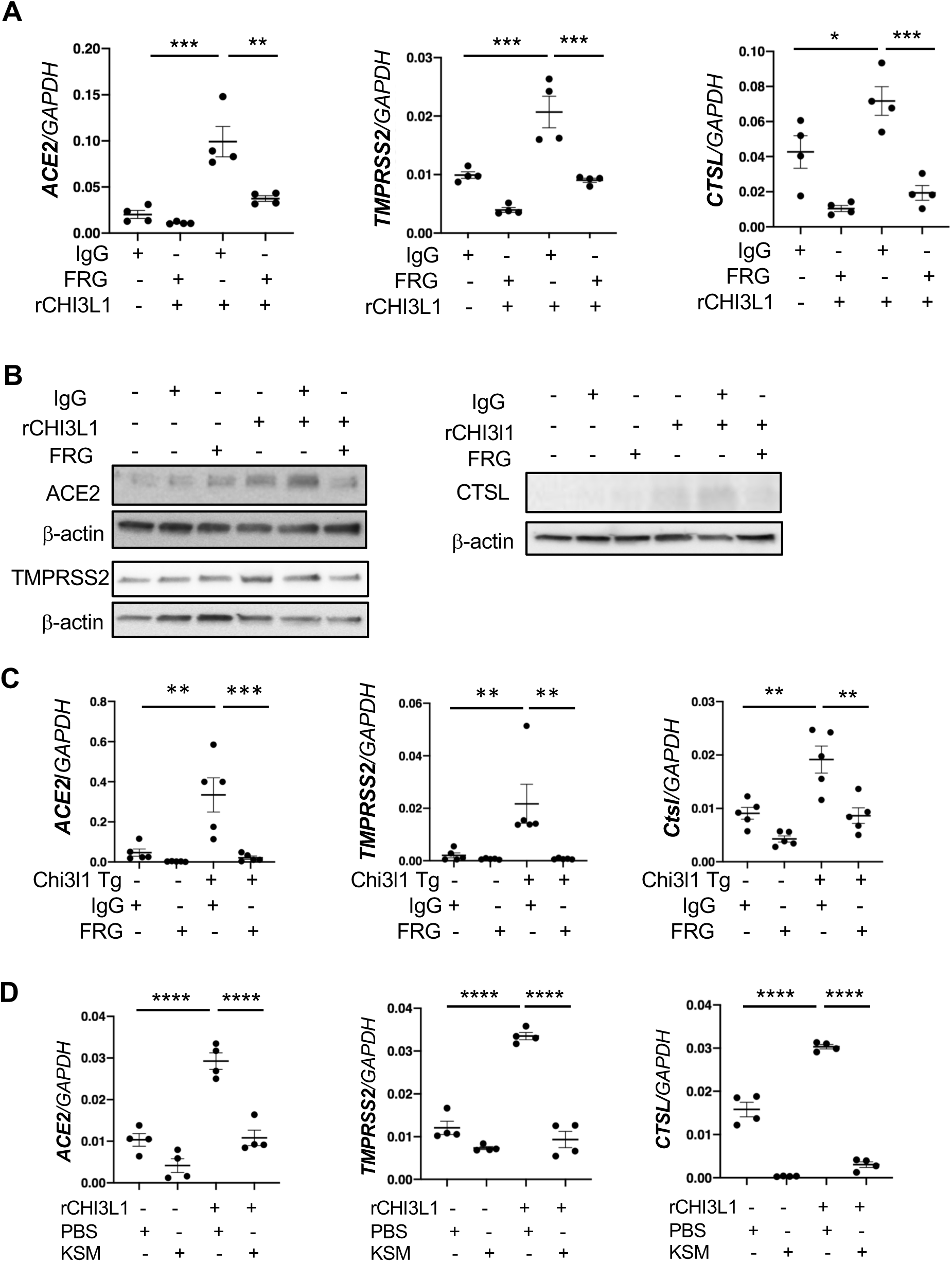
Effects of a monoclonal antibody and small molecule inhibitor on CHI3L1-stimulation of ACE2 and SPP *in vitro* and *in vivo*. (A-B) Calu-3 cells were stimulated with vehicle (PBS) or rCHI3L1 (250ng/ml) and treated with isotype control IgG or anti-CHI3L1 (FRG) for 48 hrs. The cells were then harvested and the levels of mRNA encoding ACE2 and SPP and ACE2 and SPP protein were evaluated by RT-PCR (A) and immunoblot assays (B). (C) WT and Chi3l1 Tg (Tg) mice were treated with IgG isotype control antibody or FRG antibody during their 2 weeks of transgene activation. The levels of mRNA encoding Ace2 and SPP were then evaluated using RT-PCR. (D) Calu-3 cells were stimulated with vehicle (PBS) or rCHI3L1 (250ng/ml) and treated with kasugamycin or vehicle control (PBS) for 48 hrs. The cells were then harvested and the levels of mRNA encoding ACE2 and SPP were evaluated by RT-PCR. Each value in panels A and C-D is from a different animal and the mean±SEM are illustrated. Panel B is representative of two independent experiments. \**P*<0.05, \*\**P*<0.01, \*\*\**P*<0.001, \*\*\*\**P*<0.0001 (One-Way ANOVA with post hoc Turkey multiple comparison test).

### CHI3L1 phosphorylation-based therapeutics

To date, all studies of CHI3L1 have assumed that it is not post-translationally modified. However, phosphorylation site prediction analysis using the NetPhos program (v2.0) suggests that CHI3L1 is a phosphoprotein with high potential for serine/threonine phosphorylation. Further investigation also provided a number of lines of evidence that suggest that CHI3L1 is effective as a phosphoprotein. This included (a) sequence mining of CHI3L1 which revealed a cyclin binding domain that is highly predictive of CDK phosphorylation (Supplemental Fig. S4); (b) sites of serine phosphorylation of rCHI3L1 confirmed with immunoblot assays with an anti-phosphoserine antibody (EMD Millipore, AB1603) (Fig. 4A); (c) the demonstration that CHI3L1 phosphorylation is dependent on CDK activity based on its inhibition by the broad spectrum CDK inhibitor Flavopiridol. As a result we hypothesized that phosphorylation in this region is essential for CHI3L1 effector responses. To address this hypothesis we used *in vitro* and *in vivo* approaches. In the former, we generated WT CHI3L1 and mutant forms of rCHI3L1 that could not be phosphorylated and compared their ability to induce epithelial cell ACE2 and SPP. As can be seen in Fig. 4B, WT CHI3L1 was a powerful stimulator of epithelial ACE2 and SPP and this effect was abrogated by mutations at AA 230 (Serine to Arginine mutation at site of amino acid 230 of CHI3L1). Mutations at AA 235 or 237 did not have similar effects (data not shown). In the *in vivo* experiments, we compared the effects of transgenic CHI3L1 in mice treated with the broad spectrum CDK inhibitor Flavopiridol (Fig. 4C) or the selective CDK 1 and 2 inhibitor (BMS 265246) and their vehicle controls (Fig. 4D). In both cases, the CDK inhibitors abrogated the ability of CHI3L1 to stimulate epithelial cells ACE2 and SPP. These studies demonstrate that CHI3L1 is effective as a phosphoprotein and highlight the ability of CDK inhibitors, particularly inhibitors of CDK 1 and 2, to control this phosphorylation and, in turn, serve as a therapeutic in COVID-19.

**Fig 4.**
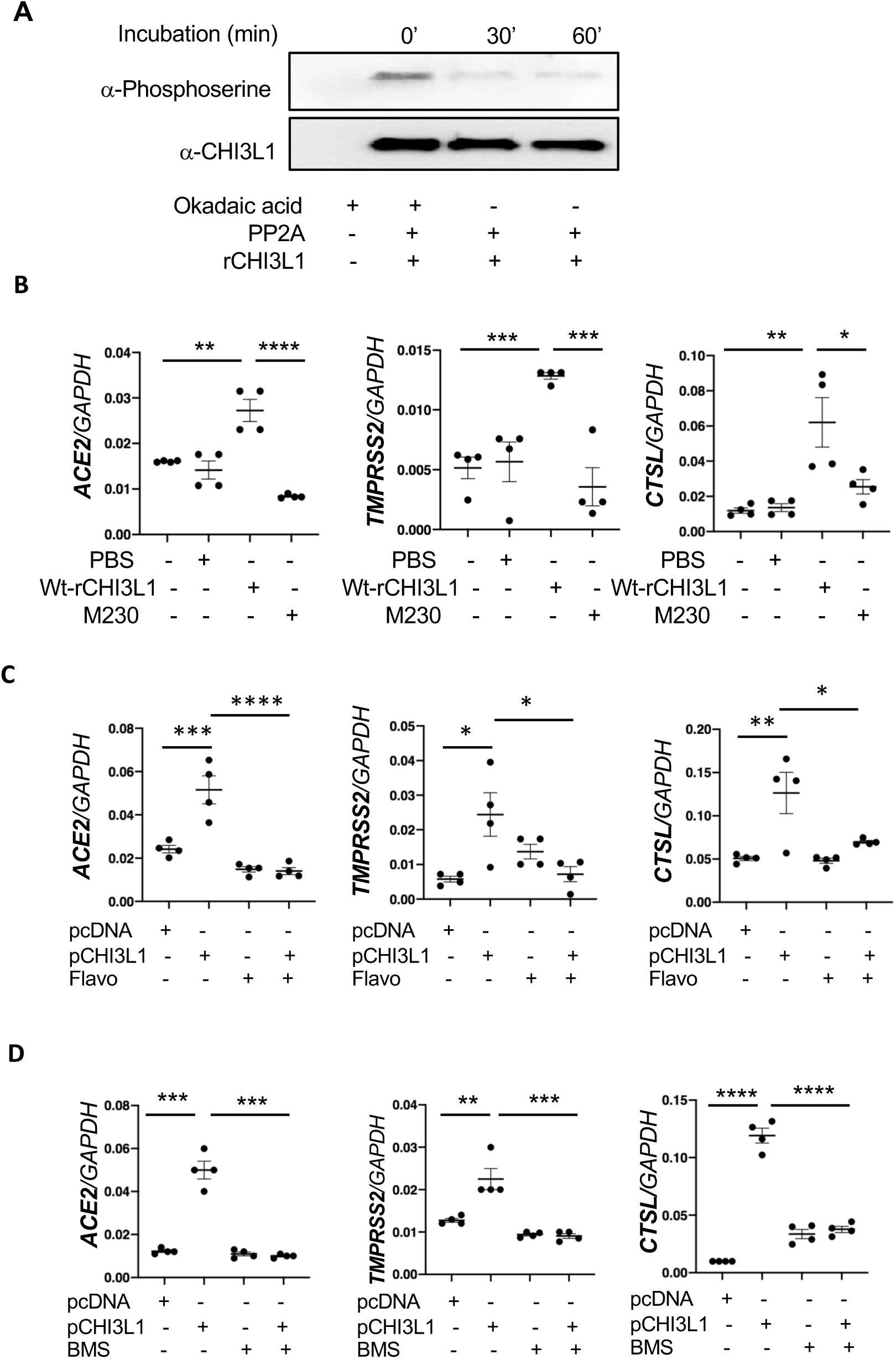
Alterations in CHI3L1 phosphorylation modify its ability to regulate ACE2 and SPP. (A) Demonstration that CHI3L1 is a phosphoprotein. Biologically active rCHI3L1 was treated with protein phosphatase 2A (PP2A) in the presence and absence of the PP2A inhibitor Okadaic acid. The alterations in CHI3L1 phosphorylation were evaluated with immunoblots using antibodies against phosphoserine or CHI3L1 controls. (B) Calu-3 cells were stimulated with a wild type (Wt) or a mutant form of rCHI3L1 (250 ng/ml for each; 24 hours) that could not be phosphorylated. The latter was done by generating a Serine to Arginine mutation at amino acid 230. The cells were then harvested and the levels of mRNA encoding ACE2 and SPP2 was evaluated by RT-PCR. (C-D) Calu-3 cells were transfected with pcDNA (vector only) or the plasmid containing a CHI3L1 cDNA (pCHI3L1). They were simultaneously treated with Flavopiridol (Flavo; 5nM) (C) or BMS265246 (BMS; 9nM) (D) or their vehicle controls (5% DMSO) for 24 hrs. The cells were then harvested and the levels of mRNA encoding ACE2 and SPP were evaluated by RT-PCR. Panel A is representative Immunoblots in 3 separate experiments. The values in panels B and D are mean±SEM. Each value in panel C is from a different animal and the mean±SEM are illustrated. \**P*<0.05, \*\**P*<0.01, \*\*\**P*<0.001, \*\*\*\**P*<0.0001 (One-Way ANOVA with post hoc Turkey multiple comparison test).

### CHI3L1 and Aging in COVID-19

The elderly is at increased risk of contracting COVID-19 and experience higher complication and case fatality rates (15, 16). These unique features have interesting parallels in the CHI3L1 axis where the levels of circulating CHI3L1 increase with aging (59). To further understand these relationships, we measured the levels of circulating CHI3L1 in WT mice as they age. As can be seen in figure 5A, the levels of circulating CHI3L1 and the levels of downstream ACE2, and SPP increased significantly in comparisons of 6 and 12-month-old WT mice. Importantly, treatment of WT mice twice a week with the monoclonal antibody FRG from 6 months to 12 months of age was remarkably effective in inhibiting the expression and accumulation of CHI3L1, ACE2 and SPP in aged mice (Fig. 5B). These studies demonstrate that the increase in circulating CHI3L1 that is seen with aging stimulates the expression and accumulation of ACE2 and SPP. They also suggest that these changes in CHI3L1, ACE2 and SPP contribute to the pathogenesis of the heightened COVID-19 responses in the elderly.

**Fig 5.**
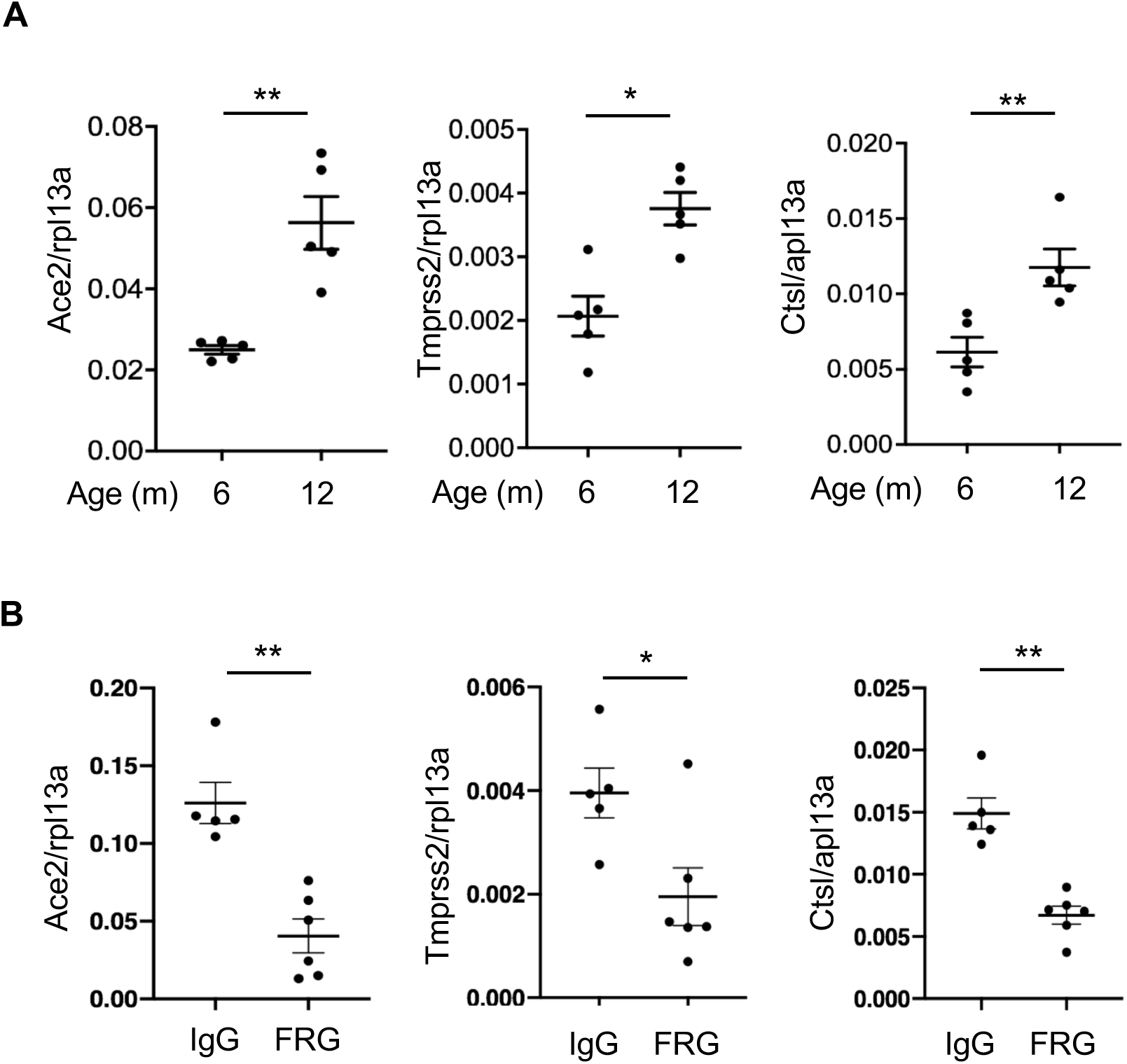
CHI3L1 is induced and regulates the expression of ACE2 and SPP in the lungs of aged mice. (A) Comparison of the levels of mRNA encoding Ace2 and SPP in lungs from 6-and 12-months old WT mice evaluated by RT-PCR. (B) Wild type mice were treated with IgG isotype antibody or anti-CHI3L1 (FRG) (200 μg/mouse, twice a week, i.p. injection) when the mice were between 6 and 12 months of age. At the end of this interval, the mice were sacrificed and the levels of mRNA encoding pulmonary Ace2 and SPP were evaluated by RT-PCR. Each value in panels A and B is from a different animal and the mean±SEM are illustrated. \**P*<0.05, \*\**P*<0.01 (Mann-Whitney U test).

### Circulating CHI3L1 in COVID-19

As noted above, SC2 infections are more common and more severe in the elderly and patients with comorbid diseases (1-3, 5, 15, 19, 20). Studies from our laboratory and others have also demonstrated that the levels of circulating CHI3L1 are also increased in the diseases that are COVID-19 risk factors (47-54). Thus, to further understand the relationships between CHI3L1 and the COVID-19 risk factors, we measured the levels of CHI3L1 in the serum of normal healthy individuals and patients presenting to the emergency department (ED) at Rhode Island Hospital for medical evaluations that prompted a COVID-19 diagnostic evaluation. The demographics of these cohorts can be seen in supplemental Table S1. In keeping with reports from our laboratory and others the majority of the normal healthy individuals had levels of circulating CHI3L1 between 15 and 60 ng/ml (Fig. 6A). Interestingly, the levels of circulating CHI3L1 in the individuals in the normal healthy cohort was not significantly different than the levels in patients presenting to the ED who did not have comorbid diseases (hypertension, diabetes, arthritis, neurologic disease, cancer, stroke, obesity and or chronic lung disease) (Fig. 6A). They were however, significantly lower than the levels in the circulation of patients presenting to the ED with existing comorbid diseases (Fig. 6A) and patients that tested positive for COVID-19 (Fig. 6B). When the COVID-19 (-) and (+) ED patients were evaluated together the levels of circulating CHI3L1 were increased in the patients that were greater than 50 years of age (Fig. 6C) and or had hypertension (Fig. 6D). The levels of circulating CHI3L1 correlated with COVID-19 disease severity. This can be seen in Figures 6E and 6F which demonstrate that the levels were significantly increased in patients that were admitted to the hospital compared those that were discharged to home (Fig. 6E) and that the circulating levels of CHI3L1 were significantly increased in patients with higher COVID Severity Scores (CSS) (Fig. 6F and Table S2). When only COVID-19 (+) patients were evaluated, the levels of circulating CHI3L1 were increased in patients with hypertension versus those without hypertension; patients that were admitted versus those that were sent home from the ED; patients that were greater than 50 years old versus younger individuals and patients with comorbid diseases versus COVID-19 (+) patients without concurrent comorbid disorders (Fig. 6, G-J). Overall, these studies demonstrate that the levels of circulating CHI3L1 are increased in the elderly and patients with comorbid disease like hypertension. They also demonstrate that the levels of circulating CHI3L1 are increased in COVID-19 (+) patients that are elderly, have comorbid diseases and manifest severe COVID-19. These observations support the concept that SC2 co-opts the CHI3L1 axis to stimulate ACE2 and SPP which augment viral infection and foster SC2 disease manifestations. They also suggest that the effects of the COVID-19 risk factors are mediated, at least in part, by their ability to stimulate CHI3L1.

**Fig 6.**
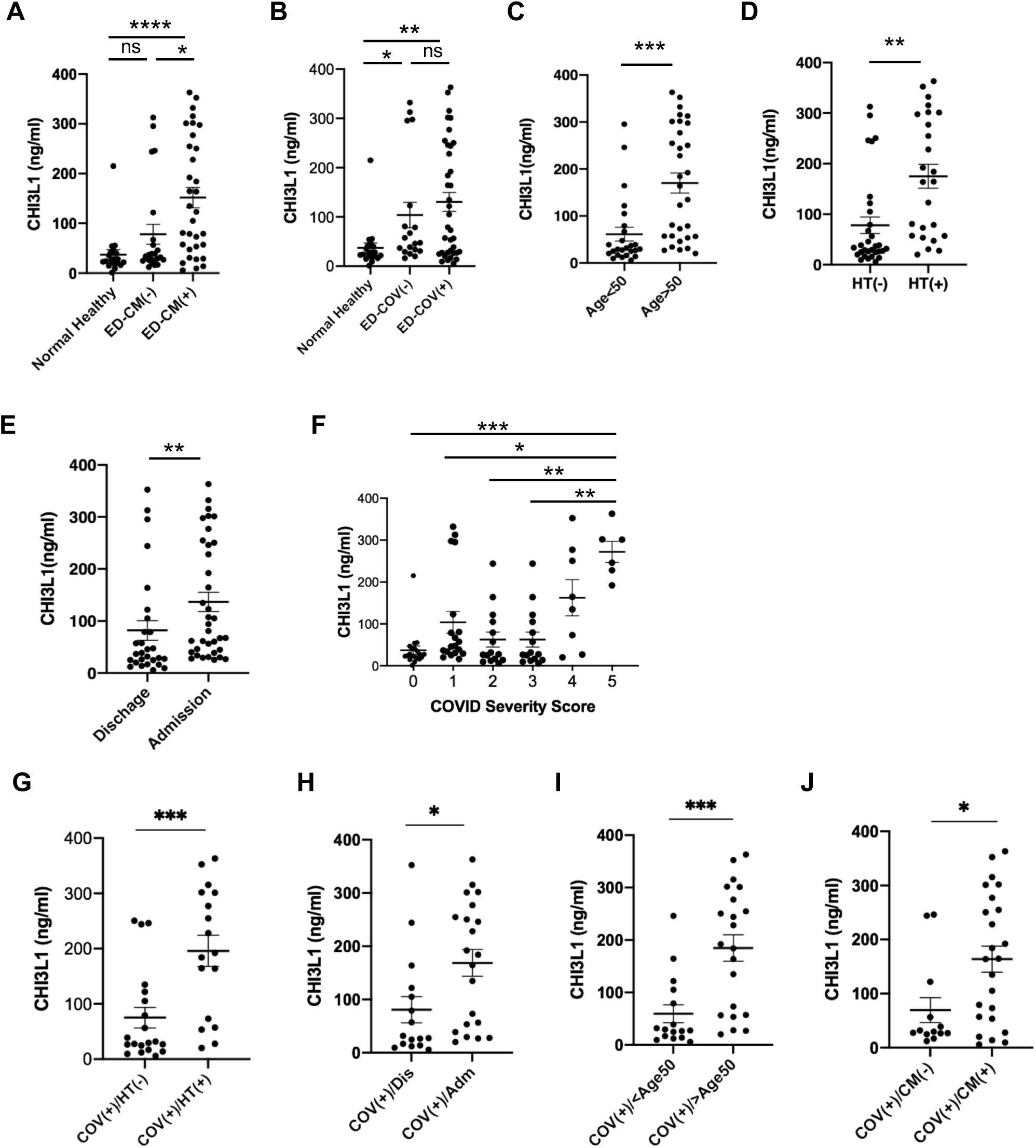
The levels of CHI3L1 in the circulation of patients with risk factors for COVID-19. The levels of CHI3L1 were assessed by ELISA using plasma from normals without comorbid diseases and patients presenting for emergency department evaluation. The values that were obtained were compared to the clinical features of the patients and the course of their diseases. ER, emergency room; CM, comorbid disease; HT, hypertension; COV, COVID-19; Dis, discharge; Adm, admission; ICU, intensive care unit. Each value is from a different individual and the mean±SEM are illustrated. \**P*<0.05, \*\**P*<0.01, \*\*\**P*<0.001, \*\*\**P*<0.0001, ns = not significant (Panels *A, B, F:* Kruskal-Walis with Dunn’s post hoc test for multiple comparisons; Panels C-E and G-J: Mann-Whitney U test). Global *P* value of panel F was 0.0002.

## Discussion

The CHI3L1 axis is a major endogenous healing and repair response and ACE2 is a protective moiety in acute lung injury (ALI) (38, 55-57). To determine if they are related to one another, we characterized the effects of CHI3L1 on ACE2 and SPP *in vivo* and *in vitro*. In our initial studies, we compared the expression of Ace2 and SPP in lungs from CC10-driven CHI3L1 overexpressing Tg mice and WT controls. These studies demonstrated induction of Ace2, Tmprss2 and CtsL in lungs from Tg (+) mice. Ace2 was most prominent in airway epithelium. It was also seen in alveoli where it co-localized with SP-C and CC10 and stained blood vessels where it co-localized on endothelial and smooth muscle cells. Tmprss2 and Ctsl were also seen in airway epithelial cells and Tmprss2 was seen in alveolar macrophages. In the *in vitro* experiments rCHI3L1 impressively stimulated the levels of mRNA encoding ACE2, TMPRSS2 and CTSL in human lung epithelial cell lines (Calu-3, A549) and primary human small airway epithelial cells (HSAEC) and lung fibroblasts. Importantly, these inductive responses were markedly decreased by treatment with moieties that alter CHI3L1-induced effector responses such as monoclonal anti-CHI3L1 (FRG), kasugamycin and inhibitors of CHI3L1 phosphorylation. These findings support the concept that SC2 co-opts the CHI3L1 healing and repair response to increase the number of cellular targets for viral entry. They also suggest that the host’s attempt to heal and repair via CHI3L1 is complicit in SC2 viral infection and COVID-19 morbidity and mortality.

In the time since the outbreak of the COVID-19 pandemic unique features of its epidemiology have been appreciated. This includes the impressive increase in the prevalence and severity of the disease in the elderly, particularly those in congregate care settings (16, 17). The prevalence and severity of COVID-19 are also increased in patients with comorbid diseases including diabetes, hypertension, obesity and metabolic syndrome, cardiovascular disease and chronic lung diseases like COPD where they associate with enhanced susceptibility to SC2 infection, more severe disease and a higher mortality (1-3, 5, 15, 19, 20). These unique features have interesting parallels in CHI3L1. Specifically, the levels of circulating CHI3L1 increase with aging. In fact, they have been reported to be the best predictor of all-cause mortality in octogenarians (59). In addition, studies from our laboratory and others have demonstrated that the levels of circulating CHI3L1 are also increased in the very diseases that are COVID-19 risk factors (47-54) where they correlate with disease severity (49, 50, 53). ACE2 expression and viral loads have been suggested to increase with aging and it has been proposed that these changes can explain the impressive severity of COVID-19 in the elderly and patients with comorbid diseases (16, 17). Our studies support this contention and provide novel insights into the mechanisms that may underlie these associations by demonstrating that the levels of circulating CHI3L1 increase during murine aging, that these inductive events are abrogated by treatment with anti-CHI3L1 and that CHI3L1 stimulates ACE2 and SPP which increases ACE2-S protein-cell interaction, S protein activation and pseudovirus-S infection.

COVID-19-associated vasculitis and vasculopathy are now considered defining features of the systemic disease caused by SC2 (10, 60). The importance of vascular alterations can also be appreciated in recent studies comparing the lungs of patients infected with SC2 and influenza (13). Both manifest diffuse alveolar damage and perivascular infiltrating lymphocytes. There were, however, distinctive vascular manifestations in the COVID-19 tissues. They included (a) severe endothelial injury with intra-endothelial SC2 virus and disrupted endothelial membranes; (b) widespread vascular thrombosis; and (c) elevated levels of intussusceptive angiogenesis. They also noted increased numbers of ACE2+ endothelial cells in infected patients versus controls (13). These and other studies have led to the belief that endothelial injury plays a key role in these responses and that the injury is caused by direct viral infection and perivascular inflammation (10, 13). Our studies support this concept by demonstrating, for the first time, that CHI3L1 is a powerful stimulator of endothelial cell and vascular smooth muscle cell ACE2 and SPP. They also raise the intriguing possibility that the CHI3L1 is a key stimulator of the pulmonary and systemic manifestations of COVID-19 and that it does this, at least in part, by enhancing the susceptibility of endothelial cells and smooth muscle cells to infection with the virus.

Studies from our lab and others have demonstrated that CHI3L1 is a critical regulator of inflammation and innate immunity and a stimulator of Type 2 immune responses, fibroproliferative repair and angiogenesis (34, 37, 38, 61-65). These studies also demonstrated that CHI3L1 is dysregulated in a variety of diseases characterized by injury, inflammation and or tissue remodeling (31-33, 37, 39-46). In keeping with these findings, we focused our recent efforts on the development of CHI3L1-based interventions that could serve as therapeutics for CHI3L1-dependent diseases. These studies have defined an exciting therapeutic platform based on CHI3L1. Our initial studies focused on biologic approaches and the generation of a panel of monoclonal antibodies against CHI3L1 using full length and peptide immunogens. These antibodies were assessed in a variety of murine models. The most impressive responses were obtained with a monoclonal antibody against a peptide that contained the sequence between AA 223-234 of human CHI3L1. This antibody is now called FRG. It was originally a mouse IgG2b kappa. A humanized version has been generated on an IgG1 backbone. There are a number of reasons to believe that FRG can be an effective therapeutic in COVID-19. First, FRG is a potent inhibitor of the basal levels and the CHI3L1-stimulated expression and accumulation of ACE2 and SPP. It is also important to point out that studies from our lab and others have demonstrated that CHI3L1 drives inflammation in a type 2 direction and augments type 2 and decreases type 1 T cell differentiation (34, 66). This is problematic for COVID-19 because, in contrast to type 2 immunity, type1 immune responses have potent antiviral effects (67). Thus, FRG treatment would abrogate CHI3L1 stimulation of type 2 responses and augment type 1 antiviral responses in a number of cells and tissues. FRG also inhibits fibroproliferative repair which could ameliorate the pathologic fibrosis that occurs in patients with respiratory failure due to COVID-19 who are intubated for an extended interval (68). Based on these observations one can see how FRG could be used as a prophylactic to diminish the chances of an individual getting infected after exposure to a SC2-infected person. One can also see how FRG, alone or in combination with antivirals such as remdisivir, could be therapeutically useful in patients with established SC2 infection.

Kasugamycin was isolated from *Streptomyces kasugaensis* in 1965 (69). It was initially appreciated to inhibit the growth of fungi, later noted to have modest antibacterial properties and most recently shown to inhibit influenza and other viral infections (70). This prompted us to undertake experiments designed to determine if Kasugamycin altered the effects of CHI3L1. These studies demonstrated that Kasugamycin is a powerful inhibitor of CHI3L1 induction of epithelial ACE2 and SPP. Other studies also demonstrated that kasugamycin inhibits type 2 immune responses and pathologic fibrosis (Elias JA, Lee, CG unpublished observation). The ability of Kasugamycin to ameliorate CHI3L1 induction of ACE2 and SPP while exerting antiviral and antifibrotic effects and augmenting Type I immune responses suggests that it is a useful agent that can be used as a prophylactic or therapeutic in COVID-19. This is an interesting prospect because Kasugmycin has been used as an antibiotic in man with minimal complications (71) and in agricultural settings where EPA evaluation has demonstrated an impressive lack of toxicity (72).

Our studies demonstrate that CHI3L1 has an activated state that is phosphorylated and that this activation is mediated by CDKs. They also demonstrate that AA 230 plays a critical role in CHI3L1 effector responses. In keeping with these findings Flavopiridol, a broad spectrum CDK inhibitor, proved to be a powerful inhibitor of CHI3L1 stimulation of ACE2 and SPP *in vitro* and *in vivo*. This is likely due, in great extent, to the inhibition of CDK 1 and or 2 because BMS 265246 had similar effects. It is also tempting to speculate that, because the monoclonal antibody FRG was generated against a site of phosphorylation, that its impressive potency is the result of its ability to function as a blocking antibody and its ability to interfere with this critical phosphorylation site. These findings suggest that Flavopiridol, BMS265246 and possibly other CDK inhibitors could be useful therapeutics in COVID-19. This could be an exciting example of drug repurposing because Flavopiridol, which is also called Alvocidib, has already received orphan drug designation from the FDA and EMA for the treatment of CLL (73).

RNA viruses often trigger the RIG like helicase (RLH) innate immune system. In this response, antiviral immunity is triggered by the dsRNA and 5’ triphosphate bearing molecules that arise in the cytosol as viral replication intermediates. These moieties are recognized by widely expressed cytoplasmic sensors from the RIG-I-like receptor family including retinoic acid-inducible gene I (RIG-I) and melanoma differentiation-associated protein 5 (mda-5) (28). The RIG-I receptor with its attached replication intermediate then interacts with mitochondrial MAVS (mitochondrial antiviral signaling molecule), the central integrator of this pathway, to triggers a type I interferon (IFN)-based antiviral response (28). The RLH pathway plays a major role in viral control and clearance. Interestingly, studies from our laboratory have also demonstrated that the RLH pathway is also a powerful inhibitor of CHI3L1 expression and accumulation (74). Based on these observations one would expect that SC2 infection and subsequent RLH activation would decrease the levels of circulating CHI3L1. This would decrease the expression of ACE2 and SPP which would decrease viral infection and spread and subsequent tissue injury. In keeping with the appreciation that viruses have evolved strategies that suppress or evade antiviral immunity (75), the levels of induction of type I IFNs are decreased in patients with COVID-19 (76, 77). We believe this decreased production of type I IFNs is a major contributor to the prevalence and severity of COVID-19 because it would increase the levels of circulating Chi3l, increase the expression and accumulation of ACE2 and SPP and augment viral infection and spread.

Early strains of SARS-CoV-2 from Wuhan China showed limited genetic diversity (78). However, genetic epidemiologic investigations conducted in late February 2020 have identified an emerging D614G mutation of the SARS-CoV-2 spike protein in viral strains from Europe (78, 79). Patients infected with D614G associated SARS-C0V-2 manifest enhanced viral loads. Pseudotyped virus with G614 mutations also manifest enhanced viral infectivity based on a spike protein that is more likely to assume an “open” configuration and bind to ACE2 with enhanced avidity (78). One can envision how S protein mutations can lead to viruses that are increasingly problematic from a therapeutic perspective. However, one can also see how CHI3L1-based therapeutics which inhibit ACE2 and SPP can inhibit SARS-CoV-2 infection regardless of the S protein sequence.

Our studies demonstrate that CHI3L1 is a major stimulator of ACE2 and SPP that enhances SC2 “S” protein-receptor binding and activation and augments SC2 infection and spread. They also led us to appreciate that the levels of circulating CHI3L1 play a major role in defining the propensity for and severity of SC2 infection and contribute to the mechanisms by which aging and comorbid diseases contribute to the pathogenesis of COVID-19. To further understand the interactions between CHI3L1 and SC2, we measured the levels of circulating CHI3L1 in a cohort of normal individuals and in a cohort of patients presenting to the ED with symptoms suggestive of COVID-19. These studies support our speculations in a number of ways. First, the levels of circulating CHI3L1 in patients in the ED cohort that were infected with SC2 and patients with comorbid diseases were significantly greater than in the normal healthy controls. In accord with what has been reported in the literature (59, 80), when the ED cohort patients were evaluated as a group, the levels of circulating CHI3L1 were increased in patients that were elderly, had hypertension, had comorbid diseases or required hospitalization. Importantly, within the COVID-19 positive ED patients the levels of circulating CHI3L1 were statistically increased in the patients that were elderly, had hypertension or other comorbid diseases and or required hospitalization. These observations support the concept that SC2 co-opts the CHI3L1 axis to augment viral infection and foster its disease manifestations. They also raise the interesting possibility that quantification of the levels of circulating CHI3L1 can be useful in assessing the severity and need for hospital admission of patients presenting with COVID-19. Additional experimentation will be required to assess this speculation and further understand the relationships between CHI3L1 and COVID-19.

## Methods

### Genetically modified mice

Lung-specific CHI3L1 overexpressing transgenic mice in which CHI3L1 was targeted to the lung with the CC10 promoter (Chi3l1 Tg) have been generated and characterized by our laboratory as previously described (34, 81). These mice were between 6-12 weeks old when used in these studies. All animals were humanely anesthetized with Ketamine/Xylazine (100mg/10mg/kg/mouse) before any intervention. The protocols that were used in these studies were evaluated and approved by the Institutional Animal Care and Use Committee (IACUC) at Brown University.

### Western blot analysis

Protein lysates from macrophages and whole mouse lungs were prepared with RIPA lysis buffer (ThermoFisher Scientific, Waltham, MA, USA) containing protease inhibitor cocktail (ThermoFisher Scientific) as per the manufacturer’s instructions. 20 to 30 µg of lysate protein was subjected to electrophoresis on a 4–15% gradient mini-Protean TGX gel (Bio-Rad, Hercules, CA, USA). It was then transferred to a membrane using a semidry method with a Trans-Blot Turbo Transfer System (Bio-Rad). The membranes were then blocked with Tris-buffered saline with Tween20 (TBST) with 5% non-fat milk for 1 hour at room temperature. After blocking, the membranes were incubated with the primary antibodies overnight at 4^0^C in TBST and 5% BSA. Th primary antibodies used in this study are: α-ACE2 (Abcam, ERR4435(2), ab108252), α-TMPRSS2 (Abcam, EPR3861, ab92323), α-CTSL (R&D Systems, AF1515). β-actin (Santa Cruz Biotechnology, Sc47778). The membranes were then washed 3 times with TBST and incubated with secondary antibodies in TBST, 5% non-fat milk for 1 hour at room temperature. After 3 additional TBST washes, Supersignal West Femto Maximum Sensitivity Substrate Solution (Thermofisher Scientific) was added to the membrane and immunoreactive bands were detected by using a ChemiDoc (Bio-Rad) imaging system.

### RNA extraction and Real-Time PCR

Total cellular RNA was obtained using TRIzol reagent (ThermoFisher Scientific) followed by RNA extraction using RNeasy Mini Kit (Qiagen, Germantown, MD) according to the manufacturer’s instructions. mRNA was measured and used for real time (RT)-PCR as described previously (34, 36). The primer sequences used in these studies are summarized in supplemental Table S3. Ct values of the test genes were normalized to internal housekeeping genes such as β-actin, GAPDH or RPL13a.

### Immunohistochemistry

Formalin-fixed paraffin embedded (FFPE) lung tissue blocks were serially sectioned at 5 μm thickness and mounted on glass slides. After deparaffinization and dehydration, heat-induced epitope retrieval was performed by boiling the samples in a steamer for 30 minutes in antigen unmasking solution (Abcam, antigen retrieval buffer, 100x citrate buffer pH:6.0). To prevent nonspecific protein binding, all sections were blocked in a ready-to-use serum free protein blocking solution (Dako/Agilent, Santa Clara, CA) for 10 minutes at room temperature. The sections were then incubated with primary antibodies (α-Ace2 (R&D Systems, AF3437), α-Tmprss2 (Abcam, ab92323), α-Ctsl (R&D Systems, AF1515), α-CC10 (Santa Cruz Biotechnology, sc-365992), α-SPC (Abcam, ab90716), α-CD31 (BD Pharmingen, 550274), α-transgelin (Abcam, ab14106)) overnight at 4°C. After three washings, fluorescence-labeled secondary antibodies were incubated for 1 hour at room temperature. The sections were then counterstained with DAPI and cover slips were added.

### Double label immunohistochemistry

Double label immunohistochemistry was employed as previously described by our lab (34)

### Generation of monoclonal antibodies against CHI3L1 (FRG)

The murine monoclonal anti-CHI3L1 antibody (FRG) was generated using peptide antigen (amino acid 223-234 of human CHI3L1) as immunogen through Abmart Inc (Berkeley Heights, NJ). This monoclonal antibody specifically detects both human and mouse CHI3L1 with high affinity (kd ≈1.1×10^−9^). HEK-293T cells were transfected with the FRG construct using Lipofectamine™ 3000 (Invitrogen, # L3000015). Supernatant was collected for 7 days and the antibody was purified using a protein A column (ThermoFisher scientific, # 89960). Ligand binding affinity and sensitivity were assessed using ELISA techniques.

### SARS-CoV-2 pseudovirus infection

Pseudotyped SARS-CoV-2 virus which has a lentiviral core expressing GFP but with the SARS-CoV-2 spike protein (expressing D614 or G614 S protein) on its envelope were obtained from COBRE Center for Stem Cells and Aging established at Brown University and Rhode Island Hospital. A plasmid expressing VSV-G protein instead of the S protein was used to generate a pantropic control lentivirus. SARS-CoV-2 pseudovirus or VSV-G lentivirus were used to spin-infect Calu-3 cells in a 12-well plate (931g for 2 hours at 30°C in the presence of 8 Dg/ml polybrene). Fluorescence microscopic images were taken 18 h after infection. Flow cytometry analysis of GFP (+) cells was carried out 48 h after infection on a BD LSRII flow cytometer and with the FlowJo software.

### Assessment of the effects of CHI3L1 on S protein-ACE2 binding and S protein processing

The effects of CHI3L1 on the binding of ACE2 and S protein were evaluated using Calu-3 cells and recombinant S protein. In brief, Calu-3 cells were incubated with vehicle (PBS) or rCHI3L1 (500ng/ml) for 24 hrs, recombinant S protein from SARS-COV2 (GeneScript, # Z03481-100) was added to the cells and further incubated for 2hrs. The cells were then harvested and subjected to co-immunoprecipitation (Co-IP) and immunoblot assays as described by our laboratory (65). The impact of CHI3L1 on the cellular processing of S-protein was also assessed using Calu-3 cells *in vitro*. In these experiments, cells were incubated with vehicle (PBS) or rCHI3L1 (500ng/ml) for 24 hrs, harvested and lysates were produced. The lysates were then incubated with recombinant S protein for 2 hrs and the un-cleaved S and cleaved S1 and S2 proteins were evaluated by Western immunoblot analysis using anti-SARS-CoV-2 Spike antibody (Proscience, #3525).

### Generation of WT and mutant forms of recombinant CHI3L1

WT CHI3L1-histidine tagged pcDNA was obtained from the MedImmune Inc (Gaithersburg, MD). The putative serine or tyrosine phosphorylation sites of CHI3L1 located between 230-239 were mutated using methods of site-directed mutagenesis. The plasmids containing WT and 4 mutant forms of CHI3L1 (M230 (Ser-Arg), M235 (Ser-Arg), M237 (Thr-Phe), M239 (Tyr-Arg)) were transfected to HEK293 cells (ATCC^®^ CRL-1573^™^) and each recombinant protein was purified using HisPur™ Ni-NTA column (Invitrogen, Cat# 88226). The specificity of recombinant protein was further validated by SDS-PAGE and Western blot evaluations.

### Biobank and healthy donor samples

Deidentified COVID-19 (+) and (-) human plasma samples were received from the Lifespan-Brown COVID-19 Biobank at Rhode Island Hospital (Providence, Rhode Island), Brown University. Normal, healthy, COVID-19 (−) samples were commercially available form Lee BioSolutions (991–58-PS-1, Lee BioSolutions, Maryland Heights, MO).

### Clinical variables and COVID Severity Score

Available deidentified clinical variables were collected from patients and from chart review during their time in the emergency department (ED). The following categorized variables were collected including chronic diseases; chronic lung disease (such as asthma, COPD or emphysema), chronic kidney disease, chronic neurologic disease (such as Parkinsons, Alzheimers, Multiple sclerosis), heart disease, high blood pressure, autoimmune diseases (such as lupus or rheumatoid arthritis), HIV/AIDS, active cancer (currently on treatment), previous stroke, overweight or obese (overweight for height >99 percentile). Symptoms were also assessed including; breathing difficulty or shortness of breath, fever, cough, change in taste or smell, rash, gastrointestinal symptoms (such as abdominal pain, vomiting, diarrhea), neurologic symptom (including stroke-like symptoms), sore throat and chest pain. A COVID-19 severity score was assigned to each individual ED patient based on their COVID-19 infection and their clinical status. This included their comorbid diseases, symptoms, discharge or admission, oxygen treatment or ICU transfer as outlined in Table S2.

### Quantification and Statistical analysis

Statistical evaluations were undertaken with SPSS or GraphPad Prism software. As appropriate, groups were compared with 2-tailed Student’s *t* test or with nonparametric Mann-Whitney U test. Values are expressed as mean ± SEM. One-way ANOVA or nonparametric Kruskal-Wallis tests were used for multiple group comparisons. Statistical significance was defined as a level of *P* < 0.05.

### Study Approval

The IRB study protocol “Pilot Study Evaluating Cytokine Profiles in COVID-19 Patient Samples” did not meet the definition of human subject research by either the Brown University or the Rhode Island Hospital IRBs. All animal procedures and experiments were conducted according to protocols approved by the Institutional Animal Care Use Committee at Brown University.

## Author Contributions

Conception and design: SK, JAE, CGL, BM, BA; Generation of experimental resources and data collection: SK, BM, CHH, BA, CML, WED, KH, OL, MJK, HJS, EM; Analysis and interpretation: SK, BA, BM, CML, CHH, JM, EM, CGL, JAE; Drafting the manuscript for important intellectual content: JAE, CGL

## Acknowledgements

This work was supported by National Institute of Health (NIH) grants U01 HL108638 (JAE), PO1 HL114501(JAE), and R01 HL115813 (CGL) from NHLBI and RO1 AG053495 (MJK) from NIA and USAMRMC W81XWH-17-1-0196 (JAE) from Department of Defense. This work was also supported by COVID-19 Research Seed Grant from Brown University (CGL and JAE). The Lentivirus Construct Core of the COBRE Center for Stem Cells and Aging was supported by a grant from the NIH (P20 GM119943 to OL). We thank Dr. Toshiyuki Okada and Professor Shigeaki Saitoh (Institute of Life Science, Kurume University, Japan) for helpful discussions on activation mechanism of CHI3L1.

