## Supplemental Tables and Figures for "Chitinase 3-like-1 is a Therapeutic Target That Mediates the Effects of Aging in COVID-19"

### Supplementary Tables

**Table S1**. Demographic features of the healthy controls and a cohort of patients used in this study

| | Healthy Control | ED Patients |
|---|---|---|
| Sex | | |
| Male | 10 | 32 |
| Female | 10 | 25 |
| Age (mean±SD)* | 48.9±15.9 | 51.2±20.25 |
| Ethnicity | | |
| White | 15 | 18 |
| Black | 0 | 7 |
| Hispanic/Latino | 3 | 21 |
| Asian or Pacific Origin | 2 | 3 |
| Other or unknown | | 7 |
| COVID-19 | | |
| Negative | 20 | 19 |
| Positive | 0 | 37 |

*, no significant difference between healthy control vs ED patients (t-test, p=0.65)

**Table S2**. COVID Severity Score

| COVID Security Score | | Sample Size |
|---|---|---|
| 0 | COVID (-) Healthy Control | 20 |
| 1 | COVID (-) ED Visit (Chronic Disease or symptoms) | 19 |
| 2 | COVID (+) ED Visit and Discharged | 16 |
| 3 | COVID (+) ED Visit and Admitted, No Oxygen | 7 |
| 4 | COVID (+) ED Visit and Admitted, Oxygen | 8 |
| 5 | COVID (+) ED Visit and Admitted to ICU | 6 |

COVID, COVID-19; ED, Emergency Department; ICU, Intensive Care Unit

**Table S3**. Sequences of RT-PCR primers used in this study

| Species | Gene | Sequence (5'to 3') |
|---|---|---|
| Mouse | Ace2-S | TCCAGACTCCGATCATCAAGC |
| | Ace2-AS | GCTCATGGTGTTCAGAATTGTGT |
| | Tmprss2-S | CAGTCTGAGCACATCTGTCCT |
| | Tmprss2-AS | CTCGGAGCATACTGAGGCA |
| | Ctsl-S | ATCAAACCTTTAGTGCAGAGTGG |
| | Ctsl-AS | CTGTATTCCCCGTTGTGTAGC |
| | β-Actin-S | GGCTGTATTCCCCTCCATCG |
| | β-Actin-AS | CCAGTTGGTAACAATGCCATGT |
| | RPL13a-S | AGGGGCAGGTTCTGGTATTG |
| | RPL13a-AS | TGTTGATGCCTTCACAGCGT |
| Human | ACE2-S | CGAAGCCGAAGACCTGTTCTA |
| | ACE2-AS | GGGCAAGTGTGGACTGTTCC |
| | TMPRSS2-S | GTCCCCACTGTCTACGAGGT |
| | TMPRSS2-AS | CAGACGACGGGGTTGGAAG |
| | CTSL-S | CTTTTGCCTGGGAATTGCCTC |
| | CTSL-AS | CATCGCCTTCCACTTGGTC |
| | β-ACTIN-S | GCCCTGAGGCACTCTTCCA |
| | β-ACTIN-AS | CGGATGTCCACGTCACACTTC |
| | GAPDH-S | GACAGTCAGCCGCATCTTCT |
| | GAPDH-AS | TTAAAAGCAGCCCTGGTGAC |

### Supplemental Figures (Fig. S1- S4)

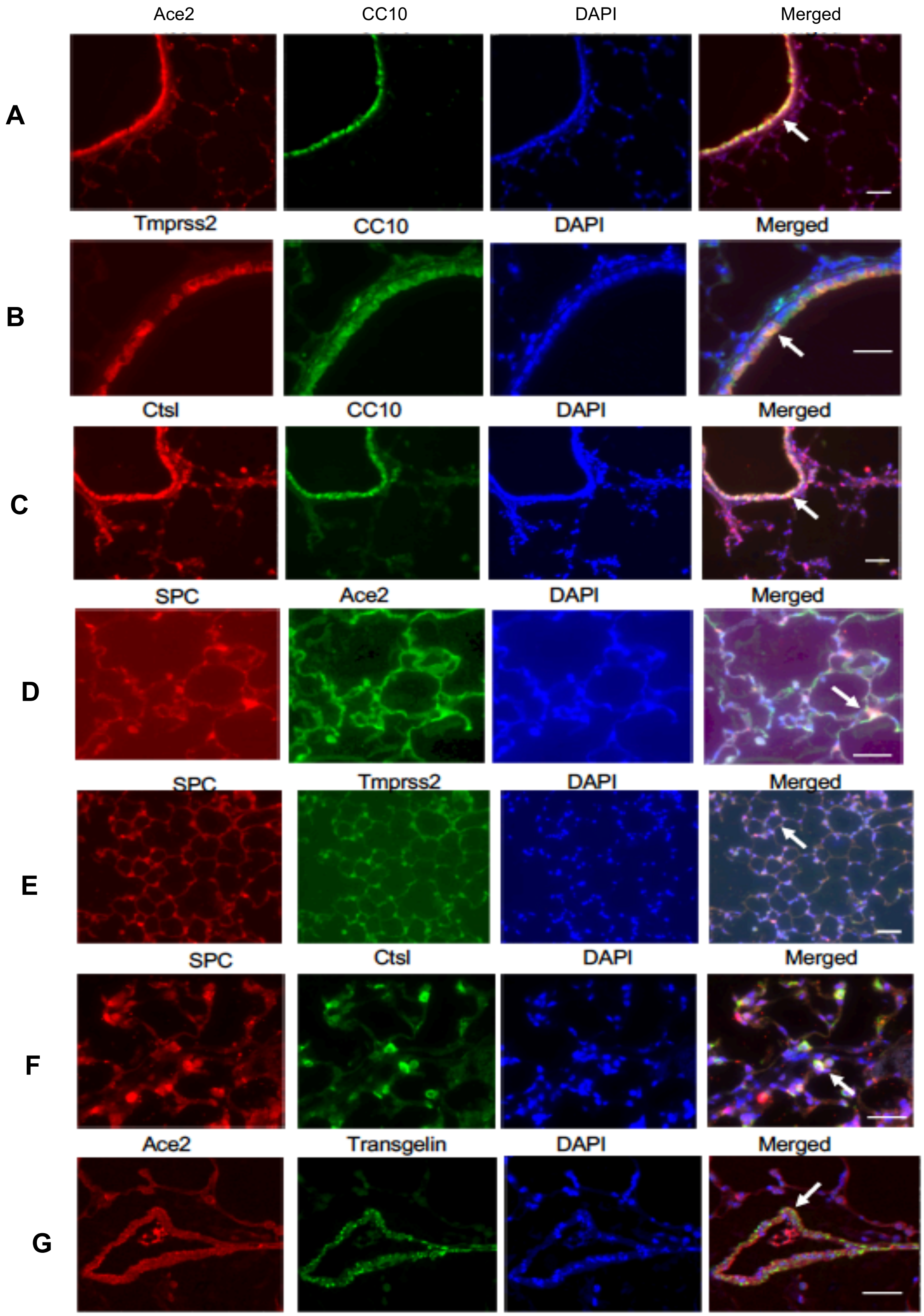

**Fig. S1. Chi3l1 induces the expression of pulmonary Ace2 and SPP**. 8 weeks old Chi3l1 Tg (+) mice were sacrificed after 2 weeks of transgene induction with Doxycycline. The cell specific expression of Ace2 and SPP in the lungs of Chi3l1 Tg mice was evaluated using double fluorescent immunohistochemistry. (A-C) Co-localization of Ace2,Tmprss2 and Ctsl with CC10, the airway epithelial cell marker in the lungs of Chi3l1 Tg mice. (D-F) Co-localization of Ace2, Tmprss2 and Ctsl with pro-SPC, the alveolar epithelial cell marker in the lungs of Chi3l1 Tg mice. (G) Co-localization of Ace2 with vascular smooth muscle cell marker Transgelin in the lungs of Chi3l1 Tg mice. Arrows in these panels indicate the stain (+) cells. Ace2, murine angiotensin converting enzyme 2; Tmprss2, transmembrane serine protease 2; Ctsl, Cathepsin L. Scale bars=100μm for panels A, C and E; Scale bars= 50 μm for panels B, D, F and G.

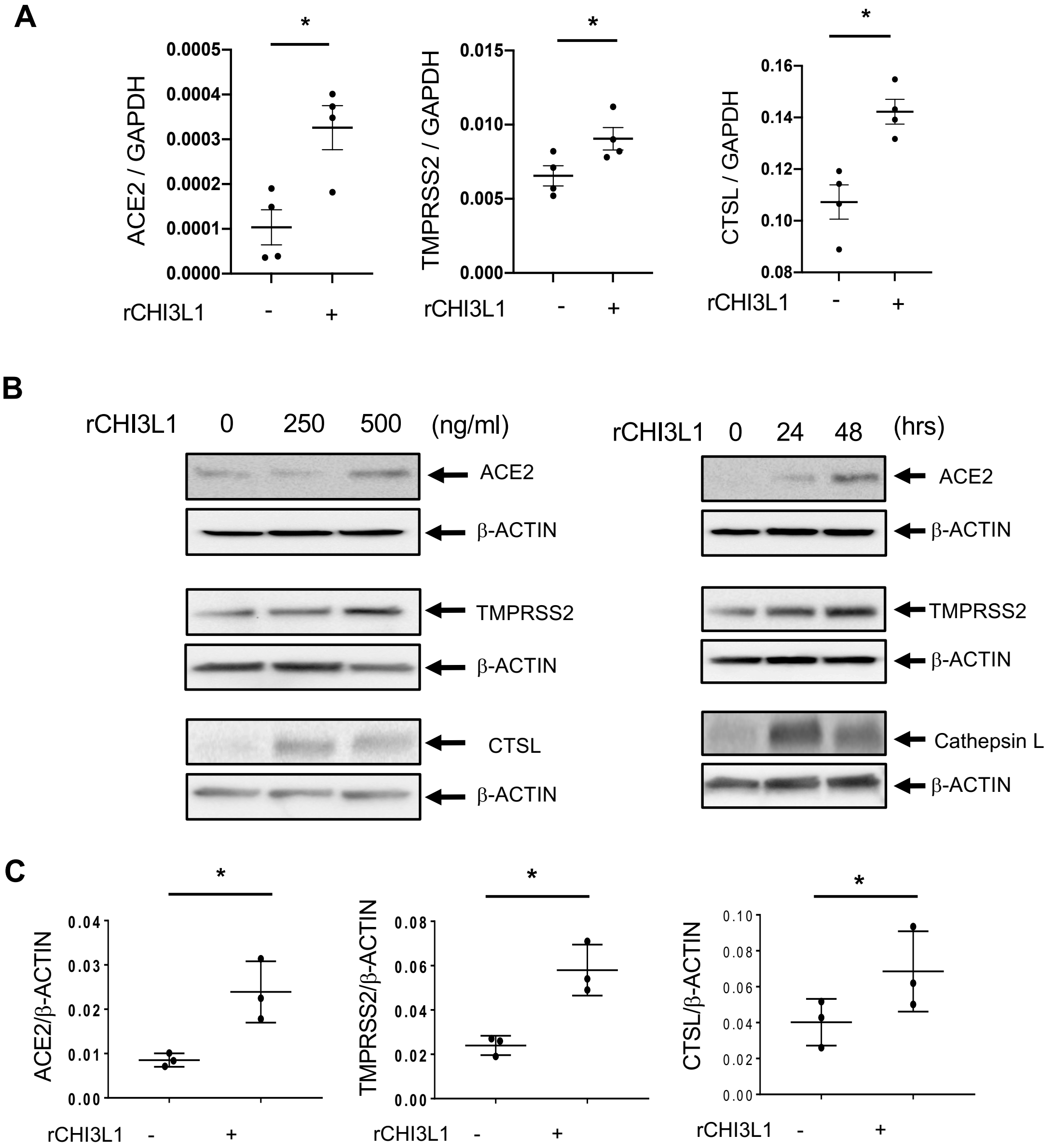


**Fig. S2**. Chi3l1 stimulates the expression of ACE2, TMPRSS2 and Cathepsin-L (CTSL) in human airway epithelial cells and fibroblasts. (A-B) A549 lung epithelial cells were subjected to real time RT-PCR (A) and Western blot evaluations (B) after stimulation of the cells with indicated dose and time. (C) After stimulation of normal human lung fibroblasts (NHLF) with CHI3L1, the expression of were stimulated. Each value in panel A and C is from a different animal and the mean±SEM are illustrated. *p<0.05, **p<0.01, ns, not significant (Mann-Whitney U test).

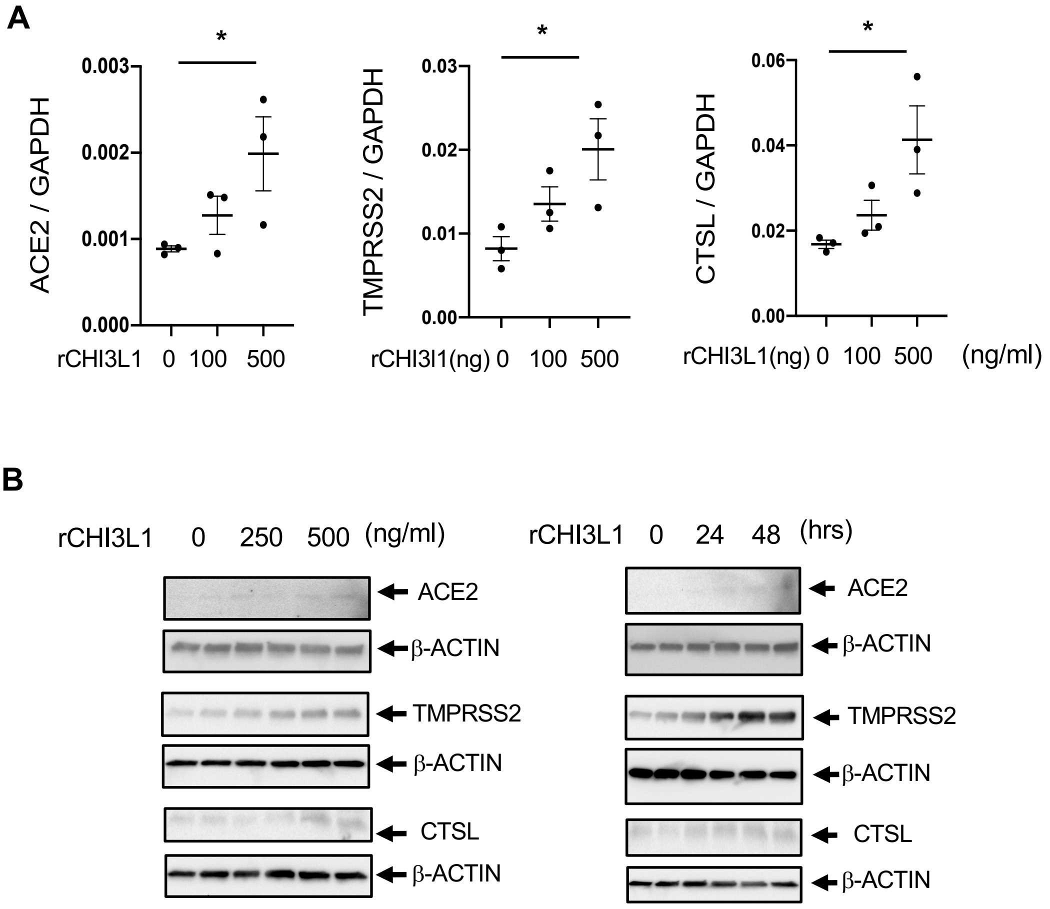


**Fig. S3**. Chi3l1 stimulates the expression of ACE2, TMPRSS2 and Cathepsin L (CTSL) in primary human airway epithelial cells. Human primary small airway epithelial cells (HSAECS) were subjected to real time RT-PCR (A) and Western blot evaluations (B) after stimulation of the cells with recombinant (r) human CHI3L1 in indicated dose and time points. Each value in panel A is from a different animal and the mean±SEM are illustrated. *p<0.05 (Kruskal-Walis with Dunn's post hoc test for multiple comparisons).

**Human CHI3L1**

MGVKASQTGFVVLVLLQCCSAYKLVCYYTSWSQYREGDGSCFPDALDRFLCTHIIYSFAN
ISNDHIDTWEWNDVTLYGMLNTLKNRNPNLKTLLSVGGWNFGSQRFSKIASNTQSRRTFI
KSVPPFLRTHGFDGLDLAWLYPGRRDKQHFTTLIKEMKAEFIKEAQPGK**KQLLL**SAALSA
GKVTIDSSYDIAKISQHLDFISIMTYDFHGAWRGTTGHH**SPL****FR**GQEDA**SPDR**FSNTDYA
VGYMLRLGAPASKLVMGIPTFGRSFTLASSETGVGAPISGPGIPGRFTKEAGTLAYYEIC
DFLRGATVHRILGQQVPYATKGNQWVGYDDQESVKSKVQYLKDRQLAGAMVWALDLDD
FQ GSFCGQDLRFPLTNAIKDALAAT

| | |
|---|---|
| **CDK phosphorylation motif** | S/T PXXK<br>S/T PXXR<br>S/T PXR |

**- Putative cyclin-binding domain**

**- Epitope for anti-Chi3l1 antibody (also called as FRG)**

**Fig.S4.** Prediction of Chi3l1 phosphorylation sites, cyclin binding domain, putative CDK activation site and illustration of the epitope used for anti-Chi3l1 antibody (FRG) generation.
